## Supplementary material for "Kinetics of blood cell differentiation during hematopoiesis revealed by quantitative long-term live imaging": Key resources table

| REAGENT or RESOURCE | SOURCE | IDENTIFIER |
| --- | --- | --- |
| Antibodies | | |
| Mouse anti-phospho-Histone H3 | Invitrogen, USA | Cat# MA3-064, RRID: AB_2633021 |
| Mouse anti-LacZ | Developmental State Hybridoma Bank, USA | Cat# 40-1a,  RRID: AB_2314509 |
| Donkey anti-mouse Cy5 | Jackson Immunoresearch laboratories Inc. | Code: 715-175-151, RRID: AB_2340820 |
| Chemicals, peptides, and recombinant proteins | | |
| VECTASHIELD with DAPI | Vector Laboratories | Cat# H-1200, RRID:AB_2336790 |
| Paraformaldehyde | ThermoFisher Scientific | Cat#28908 |
| Triton X | ThermoFisher Scientific | Cat#BP151100 |
| Normal Goat Serum | Abcam | Cat# ab7481; RRID:[AB_2716553](http://antibodyregistry.org/AB_2716553) |
| Schneider’s Drosophila medium | ThermoFisher Scientific | Cat# 21720001 |
| Fetal Bovine Serum | ThermoFisher Scientific | Cat# 12483-020 |
| Insulin solution from bovine pancreas | Sigma Aldrich | Cat# I0516 |
| Sytox Green | ThermoFisher Scientific | Cat# S7020 |
| Click-iT EdU kit | Life technologies | Cat# C10337 |
| Experimental models: Organisms/strains | | |
| *Drosophila, Tep4-Gal4>UAS-mCD8-GFP* | Gift from Dr. Lucas Waltzer, Université Clermont Auvergne, France | N/A |
| *Drosophila, dome-MESO-Gal4>UAS-mCD8-GFP* | Gift from Dr. Lucas Waltzer, Université Clermont Auvergne, France | N/A |
| *Drosophila, eater-dsRed* | Gift from Dr. Elio Sucena, Instituto Gulbenkian de Ciência, Portugal | N/A |
| *Drosophila, dome-MESO-GFP.nls* | Gift from Dr. Michele Crozatier, Université de Toulouse, France | N/A |
| *Drosophila, gstD-GFP* | Gift from Dr. Dirk Bohmann, University of Rochester Medical Center, USA | N/A |
| *Drosophila, dome-MESO-LacZ* | Gift from Dr. Nancy Fossett, University of Maryland, Baltimore, USA | N/A |
| *Drosophila, HmlΔ-dsRed.nls* | Gift from Dr. Katja Brüeckner, University of California, San Francisco, USA | N/A |
| *Drosophila, Tep4-QF>QUAS-mCherry* | Gift from Dr. Utpal Banerjee, University of California, Los Angeles, USA | N/A |
| *Drosophila, Ubi-FUCCI* | Bloomington Drosophila Stock Center, USA | RRID: BDSC_55124 |
| *Drosophila, UAS-FUCCI* | Bloomington Drosophila Stock Center, USA | RRID: BDSC_55117 |
| *Drosophila, w1118* | Tanentzapf lab, University of British Columbia, Canada | N/A |
| Software and algorithms | | |
| MATLAB | Commercial | https://www.mathworks.com/ products/matlab.html |
| FIJI | Source of the software (Schindelin et al., 2012) | https://fiji.sc/ |
| MATLAB script used to create heat maps | Codes deposited in the Tanentzapf lab GitHub (https://github.com/Tanentzapf-Lab/LiveImaging_HematopiesisKinetics_Infection_Ho_Carr) | N/A |
| MATLAB scripts used to calculate the number of prohemocytes, plasmatocyte differentiation, and total number of cells in a LG | Scripts deposited in the study (Khadilkar et al., 2017) | N/A |
| Other | | |
| Glass bottom mounting dishes | MatTek Corporation | Cat# P35G-0-14-C |
| Incubation system | TOKAI HIT | Cat# INU-ONICS F1 |
